# No support for a meiosis suppressor in *Daphnia pulex*: Comparison of linkage maps reveals normal recombination in males of obligate parthenogenetic lineages

**DOI:** 10.1101/2021.12.09.471908

**Authors:** Cécile Molinier, Thomas Lenormand, Christoph R. Haag

## Abstract

It is often assumed that obligate parthenogenesis (OP) evolves by a disruption of meiosis and recombination. One emblematic example that appears to support this view is the crustacean *Daphnia pulex*. Here, by constructing high-density linkage maps, we estimate genome-wide recombination rates in males that are occasionally produced by OP lineages, as well as in males and females of cyclical parthenogenetic (CP) lineages. The results show no significant differences in recombination rates and patterns between CP and OP males nor between CP male and CP females. The observation that recombination is not suppressed in OP males invalidates the hypothesis of a general meiosis suppressor responsible for OP. Rather, our findings suggest that in *D. pulex*, as in other species where OP evolves from CP ancestors, the CP to OP transition evolves through a re-use of the parthenogenesis pathways already present in CP and through their extension to the entire life cycle, at least in females. In addition to the implications for the evolution of OP, the genetic maps produced by this study constitute an important genomic resource for the model species *Daphnia*.

## Introduction

The mechanisms of evolutionary transitions to obligate parthenogenesis (OP) remain poorly understood. It is now clear that these transitions more often occur through modifications of meiosis rather than through replacing meiosis by mitosis (Vanin 1985; Lynch and Conery 2000; Simon *et al*. 2003). A prominent example is the small freshwater crustacean *Daphnia pulex* for which a candidate gene has been identified with a mutation that has been hypothesized to disrupt recombination in OP lineages (Hebert *et al*. 1988, 1989; Eads *et al*. 2012). Indeed, recombination is largely or entirely absent during oogenesis of OP females (Hebert and Crease 1980, 1983). However, OP lineages rarely produce males (Hebert and Crease 1983; Lynch 1984), which are genetically identical to OP females and thus carry the same mutations. These males are nevertheless known to still be able to undergo functional (*i.e*., reductional) meiosis during spermatogenesis (Innes and Hebert 1988; Xu *et al*. 2015a). However, whether or not recombination occurs during these meioses is unknown.

*Daphnia pulex* has both cyclical parthenogenetic (CP) and OP lineages, with CP being ancestral to OP. Both CP and OP share a phase of subitaneous (*i.e*., ovoviviparous) egg production, during which females parthenogenetically produce offspring whose sex is determined by the environment. The type of parthenogenesis is an aborted meiosis, which is genetically identical to mitosis (Hiruta *et al*. 2010) except for rare cases of recombination or gene conversion leading to some loss of heterozygosity (Omilian *et al*. 2006). Parthenogenetically produced males and females thus constitute a clonal lineage. CP and OP differ, however, in the mode of diapause egg production (here called diapause phase to distinguish it from the subitaneous phase). In CP, diapause egg production is sexual, whereas it is parthenogenetic in OP (Hebert 1978; Hebert and Crease 1980, 1983).

It has been suggested that, in species with CP, there may be an alternative route for the evolution of OP: OP may evolve by reusing the pathways for parthenogenetic reproduction that are already present in CP and extending them to the entire life cycle (Simon *et al*. 2003; van der Kooi and Schwander 2014). Considering these alternative mechanisms for the evolution of OP is important because transitions to obligate parthenogenesis are particularly common in species with CP (Hebert 1981; Kramer and Templeton 2001; Simon *et al*. 2002) and these species are often used as models to study the evolution of OP. Here, we study the recombination rate of rare OP males of *D. pulex*, with the aims to assess whether spermatogenetic meioses in these males involve normal levels of recombination (as compared to spermatogenetic meioses in CP males), as well as to elucidate different possible scenarios for the evolution of OP in this species.

Compared to CP males, levels of recombination in OP males might be reduced or absent for two main reasons. First, zero or very low rates of recombination in OP males may be due to the evolution of OP by a general recombination suppressor, affecting recombination during both male and female gamete formation. Indeed, suppression of meiosis or recombination is one of the main mechanisms invoked to explain transitions to obligate parthenogenesis, including in *D. pulex* (Simon *et al*. 2003; Schurko *et al*. 2009; Eads *et al*. 2012; Ye *et al*. 2021). Second, absent or reduced recombination in OP males may be due to a secondary reduction of recombination. Indeed, many forms of meiosis modifications that result in parthenogenesis do not necessarily involve recombination suppression (Bertolani and Buonagurelli 1975; Rebecchi *et al*. 2003; Oldroyd *et al*. 2008; Fougeyrollas *et al*. 2015). Yet, recombination may be deleterious under some forms of parthenogenesis, as it often leads to loss of heterozygosity, similar to inbreeding (Archetti 2004). As a consequence, there may be selection for reduced recombination within OP lineages (Engelstädter 2017). This has been documented empirically in several systems (Moritz and Haberl 1994; Rey *et al*. 2011; Boyer *et al*. 2021). Even if the meiosis modification affects female gametogenesis only, as it has been suggested for OP *D. pulex* (Innes and Hebert 1988; Paland *et al*. 2005), the secondary reduction of recombination may affect OP males as well, if it is caused by recombination modifiers that are not sex-specific.

In contrast, levels of recombination in OP males might be “normal” (meaning equal to those observed in CP) for two main reasons. First, the evolution of primary or secondary recombination suppression may involve sex-limited mechanisms, *i.e.*, involve genes that affect recombination only during female gametogenesis. Second, when OP evolved from a CP ancestor, it might have re-used the subitaneous parthenogenesis pathways already present in the ancestral CP life cycle. Specifically, in OP *Daphnia*, the parthenogenesis pathways used for subitaneous oogenesis in CP may have been simply extended to diapause oogenesis. In this case, as parthenogenesis in CP is specific to oogenesis, there may be no *a priori* reason to believe that spermatogenesis should be affected as well, which goes well in hand with the observation that OP males can achieve normal, reductional meiosis (Innes and Hebert 1988; Xu *et al*. 2015a,b).

To summarize, comparing the extent to which recombination was reduced in OP females and OP males (compared to female and male CP) can inform us on the pathways that led to OP. To date, we know that recombination is very low in OP females, but recombination rates in CP females are unknown. Indeed, recombination in CP has so far only been studied through sex-average and male-specific linkage maps (Cristescu *et al*. 2006; Xu *et al*. 2015a), but never specifically in females. Furthermore, because no previous study has addressed recombination in OP males, we do not know whether OP males recombine at a normal (CP-level) or reduced rate. In this paper, we measure these missing rates to provide a clear picture of recombination rate variation, in males and females, involved in the CP to OP transition.

As an aside, this comparison will also document the level of heterochiasmy (sex differences in recombination rates) in *D. pulex.* To date, no link between mechanisms of sex determination (genetic or environmental) and the presence of heterochiasmy has been demonstrated (Lenormand and Dutheil 2005; Stapley *et al*. 2017). However, only very few species with environmental sex determination have been studied to test any general pattern, and the data on heterochiasmy in *D. pulex* will therefore represent an interesting addition.

To compare recombination rates during diapause phase among OP males, CP males, and CP females, we performed two crosses to produce linkage maps of each of the four parents, one OP male, one CP male, and two CP females (OP females cannot be crossed and were therefore not included; their parthenogenetic recombination rate has been investigated elsewhere, though not specifically during diapause phase (Omilian *et al*. 2006; Xu *et al*. 2011; Keith *et al*. 2016; Flynn *et al*. 2017). In order to maximize the number of offspring, we used a mass mating approach with female-only clonal lines (so-called “NMP” clones for “non-male producing”). Each cross involved crossing numerous females from a CP NMP clone (a different clone in each of the two crosses) with males from another clone, either rare males from an OP clone (OP x CP cross) or males from a CP. Using Restriction-site Associated DNA sequencing (RAD-seq) we constructed highly saturated linkage maps and investigated recombination rate during gamete production in each of the four parents, according to the pseudo-testcross strategy (Grattapaglia and Sederoff 1994): SNPs that were heterozygous in both parents of a given cross (“ab × ab” SNPs) were used for the maps of both parents, while “ab × aa” and “aa × ab” SNPs (heterozygous only in the mother or only in the father) were used only for the maternal or paternal maps, respectively. For each map, meiotic recombination rates and patterns of recombination rates along the chromosomes (“recombination landscapes”) were assessed by comparing genetic and physical maps (Marey maps).

## Material & methods

### MATERIAL

We performed two mapping crosses, using four parental clones that originated from three different North American *Daphnia pulex* populations, called LPB, STM, and TEX (Table S1): The first cross, “CP x CP”, was carried out using males of the CP clone TEX-1 and females of the CP clone LPB-87, while the second cross, “OP x CP”, was carried out using rare males of the OP clone STM-2 and females of the CP clone TEX-114. Both crosses were thus inter-population crosses, and the fact that males of TEX-1 were used in one cross and females of TEX-114 in the other, allowed comparing male and female maps between clones from the same population. Both clones used as females (LPB-87 and TEX-114) are non-male producing (NMP) clones, that is, they are unable to produce males and thus they participate in sexual reproduction only as females (Innes and Dunbrack 1993; Tessier and Cáceres 2004; Galimov *et al*. 2011; Ye *et al*. 2019). The use of NMP clones meant that mass-mating could be performed without occurrence of within-clone mating (*i.e*., with obligate outcrossing between the two clones): To initiate a given cross, we introduced males of the father clone into a mass culture of the mother clone. Specifically, we regularly (about once every two weeks) introduced a small number of males into two 10L aquaria containing mass cultures of females (one for each of the two crosses), across a period of six (CP x CP) to eight (OP x CP) months. In total, 165 males were used for the CP x CP cross and 299 males for the OP x CP cross. Both crosses produced several thousands of ephippia, which were collected and stored at 4°C in the dark for two months or longer (necessary to break the diapause). Differences in male numbers used and in the duration of ephippia production were explained by the fact that many ephippia from the OP x CP cross were empty (*i.e*., did not contain any viable embryos) and because we wanted to ensure that we would be able to obtain a sufficient number of hatchlings for linkage analysis in each of the two crosses. Hatching was induced by bathing ephippia in a solution of pure water for two hours, followed by eight minutes of bleach solution and abundant rinsing with osmotic water (Retnaningdyah and Ebert 2012; Paes *et al*. 2016). The ephippia were then exposed to high light for 24h and then placed to standard laboratory conditions. The hatching vials were carefully inspected every two days for hatched juveniles, and any juvenile present was isolated individually in a new vial to initiate a clonal culture. We obtained a total of 104 clonal cultures of F1 offspring from the CP x CP cross (*i.e*., hatchlings that survived to adulthood and established a clonal culture by parthenogenesis). However, due to low hatching success, only 44 clonal cultures of F1 offspring of the CP x OP cross were obtained. All parent and offspring clones were kept under standard conditions in the laboratory, fed with the microalgae *Tetraselmis chuii*.

### DNA EXTRACTION AND RAD-SEQUENCING

One batch (offspring clones) to three batches (parent clones) of 15 to 20 individuals were collected, frozen in liquid nitrogen, and stored at -80°C. Total genomic DNA was extracted from each batch using the DNeasy® Blood & Tissue kit (Qiagen). DNA concentration and quality were examined by electrophoresis on 1 % agarose gels and with a Qubit 3.0 (high sensitivity) fluorometer. The replicate batches of the parent clones were extracted and sequenced separately to increase sequencing depth (reads from all replicates of a given parent were pooled prior to analysis). Library construction was carried out according to the RAD-sequencing protocol described by Svendsen *et al*. (2015). The libraries were sequenced on four Illumina HiSeq2500 lanes, using 100 bp single-end sequencing by the Montpellier GenomiX platform (MGX, Montpellier, France).

### SNP CALLING AND FILTERING

Raw sequencing data were demultiplexed with Stacks v.2.41 (Catchen *et al*. 2013) using process_radtags. Reads were aligned to the *D. pulex* reference genome V1.1 (Colbourne *et al*. 2011) using BWA (version: bwa-0.7.17-r1188), and reads with a mapping quality of 30 or less were removed using samtools v1.7 (Li *et al*. 2009). This procedure resulted in 5’217 to 4’695’427 reads per F1 of both crosses. Even though most F1 were well-covered (83 % of F1 had more than one million reads), also low-coverage F1 were kept because the downstream analyses in Lep-Map3 (Rastas 2017), specifically take genotypes likelihoods into account, and removal of low-coverage individuals is recommended against for these analyses. Parents were all highly covered with 3’381’813 to 6’000’981 reads per parent clone (all three replicates per parent clone combined).

The Stacks module “gstacks” with default parameters was used (--model marukilow and --var-alpha 0.05) to call SNPs and to infer genotype likelihoods. SNP markers were named according to their location, that is, scaffold name and base pair position in the reference genome. SNP markers were filtered using the module “population”, with 0.25 as the maximum proportion of missing values allowed per SNP marker across all F1 of a given cross. After this filtering step, 40’975 SNP markers were retained in the CP x CP cross and 41’917 SNP markers in the OP x CP cross.

### LINKAGE MAPS CONSTRUCTION AND ANALYSIS

#### Linkage maps

Linkage maps were constructed using Lep-MAP3 (Rastas 2017). Relationships between parents and offspring in each family were confirmed through the IBD (identity by descent) module in Lep-MAP3. The module “ParentCall2” was used to re-call missing or erroneous parental genotypes based on genotype likelihoods of the offspring, as well as to remove non-informative markers (*i.e*., markers that were homozygous in both parents). The module “Filtering2” was used to remove strongly distorted markers (*p*-value < 0.0001, as recommended for single-family data). These filtering steps reduced the numbers of retained markers to 25’951 and 32654 for the CP x CP and the OP x CP cross, respectively. The stronger reduction in the number of markers in the CP x CP cross is explained by a higher proportion of distorted markers (21 %) compared to the OP x CP cross (9 %).

The initial assignment of markers to linkage groups (LGs) followed the previous linkage map of *D. pulex* (Xu *et al*. 2015a), which was based on the same reference genome. Specifically, all markers on scaffolds that were present on the previous map, were assigned to the corresponding LGs of these scaffolds in the previous map. Second, we used the module “JoinSingles2All” to add markers on unmapped scaffolds (lodLimit=18). After the subsequent ordering steps, the initial assignment of markers to LGs was re-evaluated and corrected (if needed) using Lep-Anchor (see below). To order markers within each LG and to estimate linkage map distances, we used the module “OrderMarkers2”. The analyses were conducted separately for each parent of the two crosses using a pseudo-testcross design (Grattapaglia and Sederoff 1994).

Finally, we used Lep-Anchor (Rastas 2020) to detect potential assembly errors (“chimeric scaffold”), split them, if needed, and rerun the Lep-MAP3 pipeline using the split scaffolds. We ran three rounds of Lep-Anchor + Lep-MAP3 on the maps, until no further chimeric scaffolds were detected. This procedure identified 19 cases of likely assembly errors (assignment of parts of scaffolds to two distinct LGs or to different parts of the same LG, separated by a gap of at least 20 cM, Table S2). The final maps were based on 15’577 SNPs for LPB-87 (female of the CP x CP cross), 13’733 SNPs for TEX-1 (male of the CP x CP cross), 16’492 SNPs for TEX-114 (female of the OP x CP cross), and 21’405 SNPs for STM-2 (male of the OP x CP cross).

#### Physical distances between markers

To estimate physical distances between markers, we performed a final ordering and orientation of scaffolds, using two additional rounds of Lep-Anchor + Lep-MAP3 with SNP markers from all four linkage maps combined. This resulted in a single ordering of the scaffolds containing at least one informative marker in at least one of the four maps. Using this ordering, we estimated physical distances (in bp) between markers, using a custom R script, assuming no gaps between adjacent scaffolds and forward orientation of scaffolds whose orientation could not be determined based on the information of the linkage maps.

#### Integrated linkage map

Based on the single physical ordering of the scaffolds, we also produced a single linkage map (“integrated linkage map”) using information of both crosses. First, a sex-averaged linkage map (using the option “sexAveraged=1” in the module “OrderMarkers2”) was produced for each of the two crosses. Second, these two sex-averaged maps were combined by averaging. Specifically, for each physical position, we estimated the cM position by linear extrapolation of the nearest markers in each sex-averaged map using a custom R script and averaged these values to obtain the integrated map.

#### Recombination rate

Genome-wide recombination rate (in cM/Mb) was estimated by summing cumulative genetic lengths of all LGs and dividing it by the total length of the *D. pulex* genome, 197.3 Mb (Colbourne *et al*. 2011) or, alternatively, by the sum of the physical lengths of the anchored scaffolds 148.3 Mb. An average recombination rate for each LG was estimated using the total genetic length of a given LG, divided by the sum of the physical lengths of the scaffolds anchored on that LG.

We also estimated the within-LG recombination parameter, *r̄*_intra_ (Veller *et al*. 2019), which, in addition to the number of crossover events also takes into account their locations to estimate the average amount of shuffling of genes that occurs within a chromosome per meiosis (central and widely-spaced crossovers generate more shuffling than tightly-spaced or terminal crossovers, (Veller *et al*. 2019). To estimate *r̄*_intra_ we used the MATLAB script from Veller *et al*. (2019), considering, as measure of physical length, the total length of anchored scaffolds. Following Veller *et al*. (2019), we also estimated *r̄*_inter_, which is the probability of allele shuffling due to random assortment (*i.e.*, segregation).

### COMPARISON OF RECOMBINATION RATE BETWEEN MAPS

To visualise variation in recombination rates within LGs and to compare this variation among the different parents, we used Marey maps, which plot cumulative genetic distances (cM with respect to the first marker) against cumulative physical distances (Mb with respect to the first marker) for each marker of a given LG. The Marey maps were constructed using all markers. To quantitatively compare recombination rates between the four parents, we then used a subset of the data only (the “reduced data set”), with truncated LGs in order to ensure identical terminal positions for all four maps. Specifically, the Mb position of the most interior terminal markers on any of the four maps was used (one at each LG end), and, in maps where no marker was present at that specific physical position, the cM position was estimated by linear extrapolation of the cM positions of the two nearest markers. The cM position of all markers was subsequently adjusted (by subtracting the cM position of the first marker) to ensure that the corrected cM-position of the first marker was zero. To test for differences in total genetic lengths among the four parents, we conducted an ANOVA on genetic lengths, using each LG as a unit of replication and looking for a parental map effect. We used pair-wise post-hoc tests with the adjusted Tukey HSD method to investigate pairwise differences between any pair of parents.

To investigate potential differences in the linkage map length among the four parents at smaller scales, we divided each LG into three zones of equal length, each of them being composed of five windows (again of equal length). Linkage map positions of the boundary positions of zones and windows were estimated for each map by linear extrapolation of the linkage map positions of the nearest markers. We first tested whether specific LGs showed differences in genetic length among the parents. Second, we investigated whether specific zones within LGs showed such differences. Finally, to test for differences in crossover occurrences, independently of the total map length of the LG, we normalized all four maps to the same genetic length. Using this normalized data set, we again tested for differences among the four parents restricted to specific LGs or specific zones within LGs. Due to the frequent occurrence of windows without any crossover events, the assumptions of ANOVA were not met. We thus analysed the truncated data (both normalized and non-normalized) with pairwise, non-parametric tests: For each LG or each zone, we performed a Wilcoxon rank test (ZIW) modified for zero-inflated data (Wang *et al*. 2021). To test for effects of specific LGs, 72 (12 LGs*6 pairs) pairwise tests were performed. The effect of specific zones within LG was assessed through 216 (12 LGs*3 zones*6 pairs) pairwise tests, using windows as units of replication. The *p*-values were adjusted according to the Benjamini and Hochberg (1995) correction for multiple tests. Note that ten zones were not tested because all windows within these zones showed zero recombination in the two maps that were compared.

## Results

### LINKAGE MAPS CONSTRUCTION AND ANALYSIS

#### Linkage maps

The linkage maps of all four parent maps, including the OP male, where highly similar (Figure 1, Table 1). We therefore first present the characteristics of the integrated map only (Figure S1, Table 2). The corresponding results for the four individual maps are given in Table S3. In the second part of the results section, we then use the reduced data set to analyse potential differences among the four individual maps.

**Figure 1:**
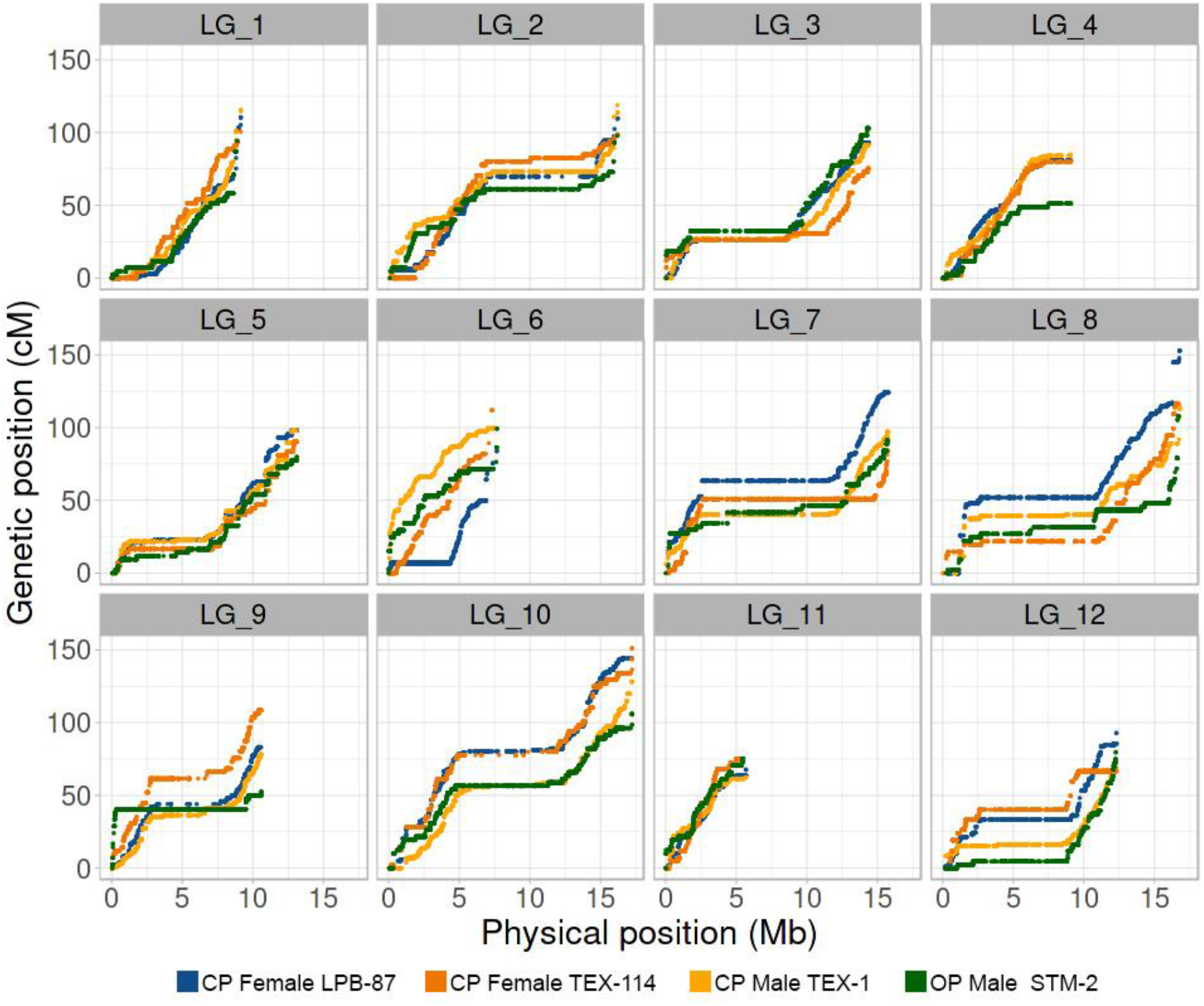
Marey maps, showing the genetic position (in cM) vs. the physical position (in Mb) of each SNP marker (dot) per linkage group (LG) and parents (color code: blue, CP_Female_LPB- 87; orange, CP_Female_TEX-114; yellow, CP_Male_TEX-1; green, OP_Male _STM-2; total, non-reduced data set in all cases).

**Table 1:** Total genetic length (in cM), total physical length of all anchored scaffolds (in Mb), and recombination rate (cM/Mb) across all LGs for each of the four parents, based on the non-reduced data set.

| Parent | Genetic length (cM) | Physical length (Mb) | Recombination rate (cM/Mb) |
| --- | --- | --- | --- |
| CP_Female_LPB-87 | 1240.60 | 142.80 | 8.69 |
| CP_Female_TEX-114 | 1160.55 | 142.22 | 8.16 |
| CP_Male_TEX-1 | 1157.16 | 144.29 | 8.02 |
| OP_Male_STM-2 | 1037.20 | 145.56 | 7.13 |

**Table 2:** Number of markers, total genetic length (in cM), total physical length of all anchored scaffolds (in Mb), and recombination rate (cM/Mb) for each LG of the integrated D. pulex linkage map. The last row (“Total”) refers to sums across all LGs, except for recombination rate where it refers to the average.

| LG | Number of markers | Genetic length (cM) | Physical length (Mb) | Recombination rate (cM/Mb) | $\bar{r}_{intra}$ |
| --- | --- | --- | --- | --- | --- |
| 1 | 3900 | 104.09 | 9.22 | 11.28 | 9.08E-04 |
| 2 | 4894 | 117.00 | 16.21 | 7.22 | 2.40E-03 |
| 3 | 3835 | 89.59 | 14.37 | 6.23 | 1.50E-03 |
| 4 | 3375 | 69.96 | 9.13 | 7.66 | 8.28E-04 |
| 5 | 3853 | 89.78 | 13.16 | 6.82 | 1.50E-03 |
| 6 | 3500 | 99.75 | 7.78 | 12.83 | 5.73E-04 |
| 7 | 4759 | 104.73 | 15.77 | 6.64 | 1.50E-03 |
| 8 | 5081 | 125.17 | 16.78 | 7.46 | 2.20E-03 |
| 9 | 4349 | 82.35 | 10.62 | 7.76 | 7.00E-04 |
| 10 | 5171 | 129.47 | 17.25 | 7.51 | 3.00E-03 |
| 11 | 2396 | 72.76 | 5.70 | 12.76 | 2.99E-04 |
| 12 | 3502 | 89.40 | 12.31 | 7.26 | 9.66E-04 |
| Total | 48615 | 1174.04 | 148.31 | 7.92 | 1.64E-02 |

#### The integrated map

The integrated *D. pulex* map contains 345 of the 5191 scaffolds of the Xu *et al*. (2015a) assembly (Table S2). Note that the LG numbering is equivalent to the one in Xu *et al*. (2015a), but we added suffixes “_1”, “_2” or “_3” for scaffolds that were split during the Lep-Anchor analysis (*i.e*., due to evidence that these likely are chimeric scaffolds). The total length of the 345 anchored scaffolds is 148.3 Mb (Table 2 and S2), which represents 75.2 % of the combined length of all scaffolds of the reference genome used here (Colbourne *et al*. 2011). The total estimated physical length of each LG ranged from 5.7 Mb on LG 11 to 17.2 Mb on LG 10 (Table 2). The four individual maps were on average 3.1 % shorter than the integrated map, missing, on average, 50 (range 41 to 64), mostly smaller scaffolds (Table S3, File S1). Our integrated *D. pulex* linkage contains a total of 48’615 SNP markers (Table 2), with an average inter-marker distance of 0.02 cM (Table 2). The total map length is 1’174 cM with the different LGs spanning between 69.96 cM on LG 4 and 129.47 cM on LG 10 (Table 2). The two sex-averaged Marey maps of each cross as well as the integrated Marey map are represented in Figure S1.

#### Recombination rates

The estimated genome-wide recombination rate of the integrated map is 7.92 cM/Mb or 5.95 cM/Mb (ranging from 5.26 to 6.29 cM/Mb among the four linkage maps), depending on whether the total linkage map length was divided by the total length of anchored scaffolds or by the estimated total genome size (197.3 Mb) of *D. pulex* (Table 2). The genome-wide intra-chromosomal recombination parameter *r̄*_intra_ across all LGs is 0.0164, while inter-chromosomal recombination parameter *r̄*_inter_ is 0.45. Recombination rates of individual LGs varied between 6.2 cM/Mb on LG 3 and 12.8 cM/Mb on LG 6 (Table 2, Figure S1), and the intra-chromosomal recombination parameter *r̄*_intra_ ranged between 3×10^−4^ on LG 11 and 3×10^−3^ on LG 10 (Table 2). The *r̄*_intra_ was positively correlated with the total genetic length (in cM) across LGs (Pearson r = 0.83, d.f. = 10, *p* = 0.0007) but negatively correlated with the recombination rate (in cM/Mb) (Spearman *ρ* = -0.68, d.f. = 10, *p* = 0.01). As evident from the Marey maps (Fig. 1), recombination rate varied extensively within LGs. In most LGs, we detected a large region with zero or almost zero recombination, putatively the peri-centromeric regions (Svendsen *et al*. 2015), although centromere locations are unknown in *D. pulex*. In contrast, recombination rates were high towards the ends of the LGs (Figures 1 and S1)

### COMPARISON OF RECOMBINATION RATE AMONG MAPS

#### Overall genetic length

All comparisons between maps were based on the reduced data set (truncated to identical terminal positions), which was 2.3 % shorter (in terms of the number of base pairs included) than the integrated map (Table S4). Overall, we found a slight but significant variation in the total genetic length among the four maps (ANOVA, *F* = 3.59, *p* = 0.02, Table 3), with only one of the pairwise post-hoc tests being significant (OP male vs. CP female LPB-87, *p* = 0.01, Table 3). Regarding sex-differences, the map length of the CP male (TEX-1) was slightly (average 6 %) but non-significantly lower than the map lengths of the two CP females (Fig. 2, Table S4, Table 3). Regarding the difference between CP and OP, the genetic length of the OP male was 11.9 % lower than that of the CP male and 15.5 % lower compared to the mean of the three CP parents (Table S4, Fig. 2). As stated above, only one of the pairwise post-hoc tests was significant (Table 3).

**Figure 2:**
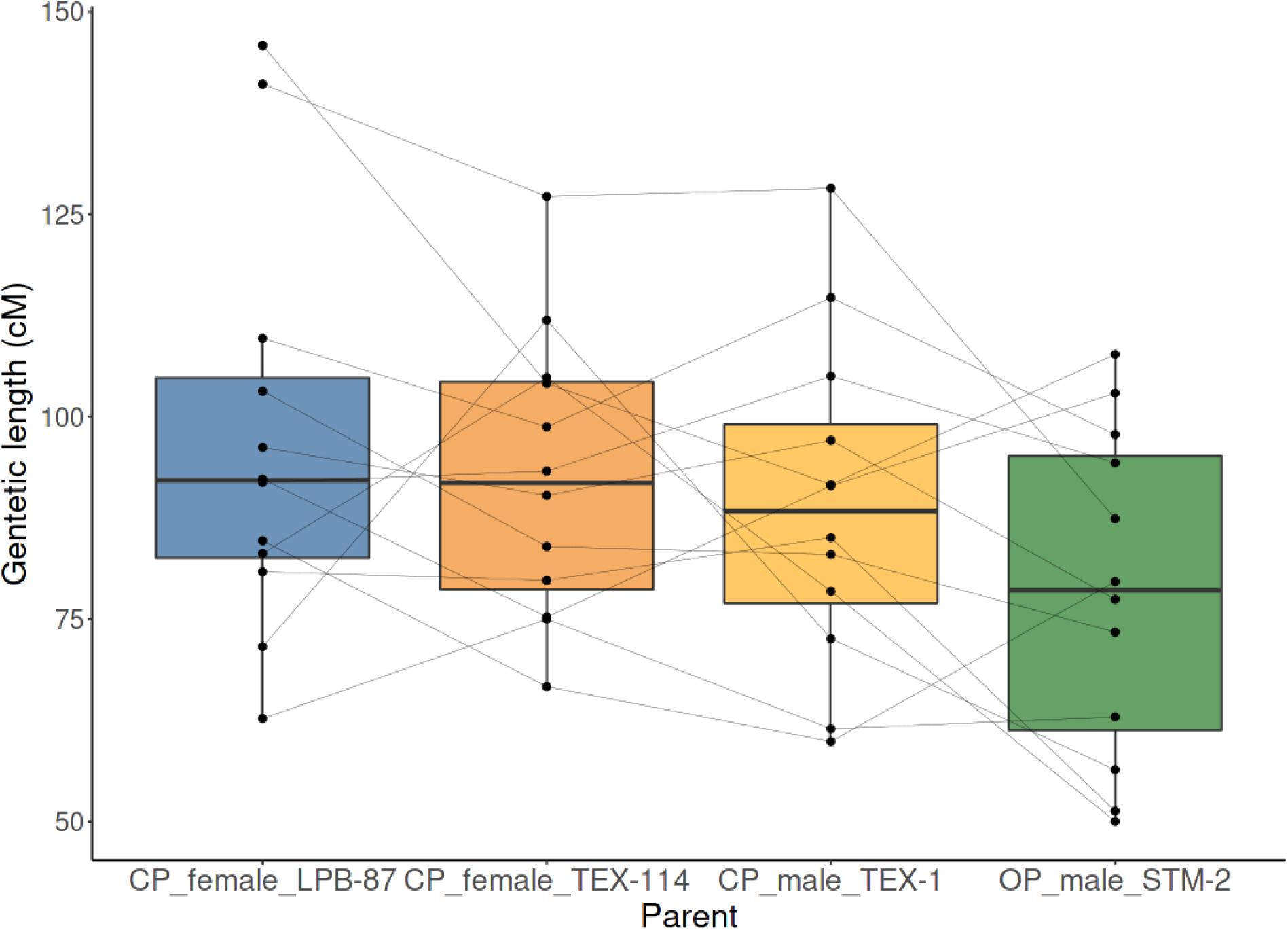
Genetic length of LGs in each of the four maps, based on the reduced data set. Dots represent individual LGs, and the fine lines identify the same LGs in the different maps. The thick horizontal lines represent the medians, the box the 25th and 75th percentiles, and the error bars are the 95 % confidence intervals. Color code as in Figure 1.

**Table 3:** Post-hoc tests for differences in the overall genetic length (using LGs as replicates) in all pair-wise comparisons between parents (“Contrast”). *P*-values are adjusted according to the Holm method.

| Contrast | **z_value** | **P_adj** |
| --- | --- | --- |
| CP_Female_TEX-114 vs. CP_Female_LPB-87 | -0.74 | 0.92 |
| CP_Male_TEX-1 vs. CP_Female_LPB-87 | -1.34 | 0.54 |
| OP_Male_STM-2 vs. CP_Female_LPB-87 | -3.14 | 0.01 |
| CP_Male_TEX-1 vs. CP_Female_TEX-114 | -0.60 | 0.92 |
| OP_Male_STM-2 vs. CP_Female_TEX-114 | -2.40 | 0.08 |
| OP_Male_STM-2 vs. CP_Male_TEX-1 | -1.80 | 0.29 |

#### Genetic length of specific LGs and zones within LGs

We tested whether the differences in total genetic length among the four maps were driven by just some of the LGs or even more narrowly by just some zones within LGs. None of the LGs differed significantly among maps (after correcting for multiple testing) in any of the pairwise comparisons (Table S5). Only LG 9 showed a tendency for being shorter (in terms of genetic length) in the OP male, compared to each of the three CP individuals (Figures 1 and 2, Table S5). Two zones within LGs showed significantly different genetic lengths among maps (Table S5): the middle zone of the LG 7 was significantly longer (*p*_adj < 0.003) in the OP male compared to each of the three CP individuals, and the middle zone of the LG 9 showed significant differences (*p*_adj < 0.003) between most pairs, being shorter in the OP male than in most CP individuals (Table S5).

#### Normalized maps

We used the normalized data set to test for differences in the localisation of crossovers, independent of the total length of the maps. Again, none of the LGs showed a significant difference in any of the pairwise comparisons and the only two zones that showed significant differences were the same ones already identified when considering non-normalized maps (Table S5).

## Discussion

### NO RECOMBINATION DIFFERENCES BETWEEN OP MALES AND CP MALE

The main goal of this study was to examine how recombination changed in males and females in the CP to OP transition. Our results demonstrate that recombination is not absent in OP males. Rather, the OP male showed highly similar levels of recombination compared to both the CP male and the CP females. While recombination rate was slightly lower than in CP, this effect was mainly local, being largely explained by LG 9 and a few zones within other LGs. These may correspond to regions that affect asexuality itself (Lynch *et al*. 2008; Eads *et al*. 2012; Tucker *et al*. 2013; Xu *et al*. 2013). The asexuality-determining regions are highly heterozygous in OP, due to hybrid origin of these regions (Xu *et al*. 2015b). This high heterozygosity (*i.e*., high levels of divergence between homologs) may be the cause of these local reductions in recombination, as demonstrated in other systems (Lukacsovich and Waldman 1999). Overall, our results clearly support the fact that OP males can be fully functional, producing sperm by a normal meiosis including normal recombination.

This contrasts with OP females, in which the diapause phase is clonal (or nearly clonal), based on the non-segregation of allozymes (Hebert and Crease 1980; Innes and Hebert 1988; Hebert *et al*. 1989). Similarly, recombination is absent (or extremely low) during the subitaneous phase of CP and OP females (Omilian *et al*. 2006; Xu *et al*. 2011; Keith *et al*. 2016; Flynn *et al*. 2017). Overall, the presence of recombination in OP males but not in OP females (diapause phase) shows that recombination suppression only concerned females, but not males in the CP to OP transition.

### POSSIBLE MECHANISM UNDERLYING THE EVOLUTION OF OP IN *D. PULEX*

#### The meiosis suppression and the Rec8 hypothesis

Given that recombination is not suppressed in OP males, it is unlikely that OP has evolved due to a *de novo* mutation leading to general suppression of recombination. General meiosis suppression, for instance due to pseudogenization (Li *et al*. 1981) of an essential recombination gene, has been put forward as one of the possible mechanisms of OP evolution (Simon *et al*. 2003; Schurko and Logsdon 2008). In *D. pulex*, a particular haplotype containing a frameshift mutation in one of the three genomic copies of the Rec8 gene (Rec8-B) consistently occurs (in heterozygous form) in OP but not in CP (Eads *et al*. 2012). Rec8 is involved in the cohesin complex that binds sister chromatids during meiosis and is therefore a good candidate for a gene that might lead to recombination suppression if its function is disrupted. Rec-8 is not specific to the female meiosis: all Rec-8 paralogs are expressed in both sexes of CP *D. pulex* (Schurko *et al*. 2009) and there is so far no evidence for male-biased or female-biased expression of Rec8-B.

However, our data indicates that disrupting Rec8-B does not lead to recombination suppression in OP males. The males in our experiments are genetically identical to OP females and therefore also heterozygous for the loss of function mutation in Rec8-B, while still having a functional copy of Rec8-B, just as the females. Thus, our result shows that there is no evidence for a causal involvement of Rec8-B in the evolution of OP. Rather, the Rec8-B mutation may have occurred secondarily in OP. Loss of function mutations can indeed occur secondarily in genes that are no longer under strong selection pressure (Normark *et al*. 2003).

#### The sex-limited meiosis suppression hypothesis

Normal recombination in OP males is consistent with a scenario where OP evolution is caused by mutation(s) affecting recombination only during oogenesis. This is the idea of a sex-limited meiosis suppression gene (Hebert *et al*. 1988, 1989). This sex-specific suppression might have occurred in CP through de novo mutations. We do not observe heterochiasmy in CP (see below), suggesting that this type of variation is not frequent, or at least those mechanisms differentially adjusting recombination in males and females do not pre-exist in CP.

Another possibility is that OP evolved by reusing the subitaneous parthenogenesis oogenesis pathways already present in CP and extending them to oogenesis during diapause egg formation. In this scenario, the sex-limited meiosis suppression is based on an already existing pathway and only requires that it becomes used in a different part of the life cycle. Because this modification is likely to be minor (*e.g.*, involve different signalling or expression patterns during diapause egg production), it may be a common route to evolve OP in *Daphnia* and other CP-OP systems. In aphids, OP has evolved though a genetic change that prevents individuals from entering the diapause phase and are typically observed in temperate regions with mild winters (Simon *et al*. 2002, 2010; Dedryver *et al*. 2013). The identified candidate region in the pea aphid (*Acyrthosiphon pisum*) contains genes involved photoperiod sensitivity (Jaquiéry *et al*. 2014). Similarly, in rotifers, the transition to OP is thought to be caused by a genetic change that prevents individuals from responding to chemical signals that induce sexual reproduction in CP (Stelzer 2008; Stelzer *et al*. 2010). In contrast to aphids, OP *Daphnia* still enter diapause phase, so that the mechanism is probably different. It cannot involve only an altered sensitivity to environmental signals. However, the general principle may be the same. Once parthenogenesis is present in a part of the life cycle, a transition to OP can simply be achieved by extending it to the entire life cycle, rather than by evolving a new, female-limited, parthenogenetic pathway.

#### Secondary evolution in OP male

In our experiment, we deliberately used an OP strain known to be able to undergo successful, reductional meiosis (Xu *et al*. 2015b). Indeed, other OP strains exist, in which males do produce diploid or aneuploid sperm (Xu *et al*. 2015b). Doing a mapping cross with a male from such a strain would either have been impossible (in case of unviability of the produced offspring) or technically too challenging (interpretation of segregation patterns in offspring with a potential mixture of diploid and triploid loci). We therefore do not know whether spermatogenesis in these males involves normal recombination. Yet, it is likely that non-reductional (and potentially non-recombining) spermatogenesis in these males is explained by secondary evolution, a scenario in line with the expected secondary loss of males or male functions in OP following a relaxation of selection pressure (Innes *et al*. 2000; Wolinska and Lively 2008; van der Kooi and Schwander 2014). Indeed, the emergence of new OP lineages occurs through contagious asexuality where males transmit OP genes, which originated in a hybrid lineage to new lineages by mating with CP females (Innes and Hebert 1988; Crease *et al*. 1989; Hebert *et al*. 1989, 1993; Taylor and Hebert 1993; Paland *et al*. 2005; Xu *et al*. 2015a). As all known OP lineages (with the exception of high arctic ones, (Beaton and Hebert 1988; Dufresne and Hebert 1995) are diploid, the males transmitting OP genes to these lineages must have been able to undergo reductional meiosis, just as in our experiment.

### NO HETEROCHIASMY IN CP *D. PULEX*

We produced both male-specific and female-specific linkage maps of *D. pulex*, which allows us to evaluate how recombination changed in males and females in the CP to OP transition. Even though the CP male recombined slightly less than the two CP females, we found no evidence for genome-wide heterochiasmy in *D. pulex*. This is the first evidence for the absence of heterochiasmy in a species with environmental sex determination and no sex chromosomes. The result is congruent with very recent finding in *D. pulicaria*, the sister species of *D. pulex*, in which also no heterochiasmy was found (Wersebe Matthew 2021). The only other case of an ESD animal where sex-specific recombination rate was investigated, is the saltwater crocodile where there is strong heterochiasmy (Miles *et al*. 2009). Hence, our findings tend to confirm that there is no special pattern of heterochiasmy in ESD species, and no global association between mechanisms of sex determination (genetic or environmental) and the presence of heterochiasmy (Lenormand and Dutheil 2005; Stapley *et al*. 2017). We also observed that the female LPB-87 has a non-recombining region at the beginning of the LG 6 but this difference was not shared with the female TEX-114, and thus is more likely to be explained by a population difference rather than by the sex. This also highlights the fact that taking into account inter-population variability may be important when studying heterochiasmy, either by using within-sex biological replicates from different populations or males and females from the same populations (both were done here).

### A NEW REFERENCE MAP FOR *D. PULEX*

The sex-specific and integrated maps presented in the current study constitutes an important addition to existing genomic resources for *D. pulex*. The first *D. pulex* linkage was based on microsatellite data (Cristescu *et al*. 2006). Subsequently, Xu *et al*. (2015a) produced a second-generation, male-specific map, based on single sperm methodology. An additional map, which was published as an annex of a new reference genome for the species (Ye *et al*. 2017), is likely erroneous, as it predicts, on average, over eight crossovers per chromosome and meiosis, as opposed to just a bit over two in our map and that of Xu *et al*. (2015a). We therefore compare our results, mainly to the linkage map from Xu *et al*. (2015a), which was also based on the same genome assembly as used here (Colbourne *et al*. 2011). Xu *et al*. (2015a) anchored 187 scaffolds (131.9 Mb) and have an average inter-marker distance of 0.87 cM, while our integrated map anchors 345 scaffolds (148.3 Mb) with 0.02 cM 0.02 cM between markers on average. The main improvement thus comes from the mapping of many additional, mostly smaller scaffolds. In addition, while there was a high degree of collinearity between the maps, we identified and corrected 19 likely assembly errors (chimeric scaffolds) and placed the part-scaffolds back to the linkage map. Still, about one fourth of the total assembly (197.3 Mb) remains unmapped, either due to smaller scaffolds containing no SNPs, scaffolds with SNPs only in repetitive regions (which are filtered during mapping due to a low mapping score), and perhaps also due to the presence of contaminant scaffolds (*e.g*., DNA from microbial symbionts) in the reference genome.

Regarding the genome-wide recombination rate, the estimates from our study and that of Xu *et al*. (2015a) are very similar (7.9 cM/Mb and 7.3 cM/Mb, respectively). These estimates are also similar to those from other *Daphnia* species (*D. pulicaria*, 7.4 cM/Mb, (Wersebe Matthew 2021) and *D. magna*, 6.8 cM/Mb, Dukić *et al*. (2016), suggesting conservation of recombination rates in the genus.

Regarding the individual maps, there appears to be large variation among individuals in the ranking of the longest to the shortest LG. Inspection of the Marey maps (Fig. 1) suggests that the differences are largely due to a small group of terminal markers per LG, while the recombination patterns were otherwise (apart from the few notable exceptions discussed above) remarkably similar among individuals. Two factors may have contributed do differences in estimated recombination rates in terminal markers among individuals. First, the observation may be entirely artefactual because the estimation of recombination rate is less reliable for terminal markers than for more central ones. Indeed, to counter the well-known fact that erroneous genotype information artificially increases recombination rate, Lep-MAP3 (Rastas 2017) uses information on several flanking markers to smoothen spikes in apparent recombination rates due to unreliable markers. Second, as most LGs exhibited higher recombination rates in more peripheral parts, the estimated total length of LGs may be rather sensitive to inclusion or not of an additional, slightly more terminal marker as well as to sampling variation among the different maps.

The high prevalence of peripheral crossovers likely has also contributed to the observed low *r̄*_intra_ (within-LG recombination parameter) because terminal recombination contributes only little to effective gene shuffling. The excess of recombination in peripheral parts was mainly noted in (physically) larger LGs, a pattern also observed in many other animal and plant species (Haenel *et al*. 2018). This pattern might amplify the very well-known negative relationship between the recombination rate (cM/Mb) and the physical size of LGs, caused by the constraint of at least one crossover per LG and meiosis (Mather 1938; Hunter 2007). It might thus also contribute to the observed positive and negative correlations of *r̄*_intra_ with cM length and cM/Mb recombination rate across LGs, which are likely explained by the same factors.

Overall, we found that the inter-chromosomal recombination parameter *r̄*_inter_ was much larger than the intra-chromosomal one, *r̄*_intra_. This is not surprising given that the species has 12 different chromosome pairs of more or less similar physical size (suggesting that the probability of a random pair of genes to be on two different chromosomes is about 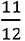), and given that recombination within chromosomes is not free. Nonetheless, this finding illustrates that the reduction of crossover numbers or an evolution to more terminal crossover locations would have minor effects on overall shuffling. This highlights the fact that even if recombination rates were reduced in OP males, gene shuffling reduction would be efficient only if segregation was reduced at the same time.

## Conclusion

We found that the CP to OP transition in *D. pulex* involves a considerable reduction in female recombination rate, that male recombination is not affected, and that recombination is not initially different between male and female CP. These findings favour the hypothesis that the subitaneous parthenogenetic pathway was re-used and extended to the production of diapause egg in *D. pulex*. This may be a common way to evolve obligate parthenogenesis in species with mixed sex-asex reproductive systems.

**File S1**: Excel file with five sheets, containing the integrated linkage map (sheet 1) and the four parental maps (sheets 2 to 5). In each sheet, each line corresponds to a marker (“Marker_ID”), whose name is based on the reference genome (scaffold and bp position within scaffold). For each marker, its LG and cM position are given, a well as its cumulative physical position within the LG (see materials and methods). Two additional columns indicate whether the marker is included in the “Reduced data” set, and whether it is on a “Split scaffold”.

**Table S1:** Names and origins of clones used in the study, as well as their use as mother or father line in each of the two crosses.

**Table S2:** Map Physical locations of the 345 anchored scaffolds in the integrated map. Scaffolds are named according to (Xu et al. 2015a), with suffixes “_1”, “_2”, or “_3” for scaffolds split during the analysis (due to evidence that the original scaffolds were chimeric). For each scaffold its location is indicated by the linkage group (LG) to which it is assigned, the start and end positions (in bp) of the scaffold within that LG, as well as the orientation (“up” for the same orientation as in the reference genome, “down” for the opposite, i.e., highest position first). For split scaffolds, bp positions after which they were split are indicated, based on the unsplit scaffold in original (i.e., “up”) orientation.

**Table S3:** Total genetic length (in cM), total physical length of all anchored scaffolds (in Mb), and number of markers for each LG and for each of the four parents, based on the non-reduced data set. Physical lengths differ slightly among parents because of different numbers of anchored scaffolds. Totals refer to sums across LGs.

**Table S4:** Physical and genetic lengths of each LG in each of the four parents in the reduced data set. Totals refer to sums across LGs. Details on the positions of the terminal markers are given in File S1.

**Table S5:** Zero-inflated Wilcoxon rank tests (ZIW) for differences in recombination between pairs of parents (“Contrast”) based on the normalized or non-normalized data set (“Data type) and either for specific LGs or specific zones within LGs (“Genome region”). *P*-values adjusted by the Benjamini & Hochberg method (Benjamini and Hochberg 1995).

**Figure S1:** Marey maps, showing the genetic position (in cM) vs. the physical position (in Mb) of each SNP marker (dot) per linkage group (LG) for the integrated map and the two sex-averaged maps from each cross (color code: black, integrated map; green and orange, sex-averaged maps from the CP x CP and OP x CP cross respectively; total, non-reduced data sets in all cases).

## Acknowledgements

The authors are grateful to Michael Lynch for kindly providing the *D. pulex* strains used. We thank Roula Zahab, Marie-Pierre Dubois and the GEMEX platform for help with the construction of the libraries. We thank the MGX-Montpellier GenomiX platform for the sequencing. We thank Pasi Rastas and Enrique Ortega-Abboud for helpful comments during the maps’ construction. This work was funded by grant ANR-17-CE02-0016-01, GENASEX, from the French National Research Agency. The authors declare no conflicts of interest.

## References

1. Archetti, M. 2004. Recombination and loss of complementation: A more than two-fold cost for parthenogenesis. J. Evol. Biol. 17:1084–1097.

2. Beaton, M. J., and P. D. N. Hebert. 1988. Geographical parthenogenesis and polyploidy in *Daphnia pulex*. Am. Nat. 132:837–845.

3. Benjamini, Y., and Y. Hochberg. 1995. Controlling the false discovery rate: A practical and powerful approach to multiple testing. J R Stat Soc Series B Stat Methodol 57:289–300.

4. Bertolani, R., and G. P. Buonagurelli. 1975. Osservazioni cariologiche sulla partenogenesi meiotica di *Macrobiotus dispar* (Tardigrada). Atti della Accad. Naz. dei Lincei. Rend. Ser. 8 53:782–786.

5. Boyer, L., R. Jabbour-Zahab, M. Mosna, C. R. Haag, and T. Lenormand. 2021. Not so clonal asexuals: Unraveling the secret sex life of *Artemia parthenogenetica*. Evol. Lett. 5:164–174.

6. Catchen, J., P. A. Hohenlohe, S. Bassham, A. Amores, and W. A. Cresko. 2013. Stacks: An analysis tool set for population genomics. Mol. Ecol. 22:3124–3140.

7. Colbourne, J. K., M. E. Pfrender, D. Gilbert, W. K. Thomas, A. Tucker, T. H. Oakley, S. Tokishita, A. Aerts, G. J. Arnold, M. K. Basu, D. J. Bauer, C. E. Cáceres, L. Carmel, C. Casola, J. H. Choi, J. C. Detter, Q. Dong, S. Dusheyko, B. D. Eads, T. Fröhlich, K. A. Geiler-Samerotte, D. Gerlach, P. Hatcher, S. Jogdeo, J. Krijgsveld, E. V. Kriventseva, D. Kültz, C. Laforsch, E. Lindquist, J. Lopez, J. R. Manak, J. Muller, J. Pangilinan, R. P. Patwardhan, S. Pitluck, E. J. Pritham, A. Rechtsteiner, M. Rho, I. B. Rogozin, O. Sakarya, A. Salamov, S. Schaack, H. Shapiro, Y. Shiga, C. Skalitzky, Z. Smith, A. Souvorov, W. Sung, Z. Tang, D. Tsuchiya, H. Tu, H. Vos, M. Wang, Y. I. Wolf, H. Yamagata, T. Yamada, Y. Ye, J. R. Shaw, J. Andrews, T. J. Crease, H. Tang, S. M. Lucas, H. M. Robertson, P. Bork, E. V. Koonin, E. M. Zdobnov, I. V. Grigoriev, M. Lynch, and J. L. Boore. 2011. The ecoresponsive genome of *Daphnia pulex*. Science 331:555–561.

8. Crease, T. J., D. J. Stanton, and P. D. N. Hebert. 1989. Polyphyletic origins of asexuality in *Daphnia pulex*. II. Mitochondrial-DNA variation. Evolution 43:1016–1026.

9. Cristescu, M. E. A., J. K. Colbourne, J. Radivojac, and M. Lynch. 2006. A microsatellite-based genetic linkage map of the waterflea, *Daphnia pulex*: On the prospect of crustacean genomics. Genomics 88:415–430.

10. Dedryver, C. A., J. F. Le Gallic, F. Mahéo, J. C. Simon, and F. Dedryver. 2013. The genetics of obligate parthenogenesis in an aphid species and its consequences for the maintenance of alternative reproductive modes. Heredity 110:39–45.

11. Dufresne, F., and P. D. N. Hebert. 1995. Polyploidy and clonal diversity in an arctic cladoceran. Heredity 75:45–53.

12. Dukić, M., D. Berner, M. Roesti, C. R. Haag, and D. Ebert. 2016. A high-density genetic map reveals variation in recombination rate across the genome of *Daphnia magna*. BMC Genet. 17:1–13.

13. Eads, B. D., D. Tsuchiya, J. Andrews, M. Lynch, and M. E. Zolan. 2012. The spread of a transposon insertion in Rec8 is associated with obligate asexuality in *Daphnia*. Proc. Natl. Acad. Sci. USA 109:858–863.

14. Engelstädter, J. 2017. Asexual but not clonal: Evolutionary processes in automictic populations. Genetics 206:993–1009.

15. Flynn, J. M., F. J. J. Chain, D. J. Schoen, and M. E. Cristescu. 2017. Spontaneous mutation accumulation in *Daphnia pulex* in selection-free vs. competitive environments. Mol. Biol. Evol. 34:160–173.

16. Fougeyrollas, R., K. Dolejšová, D. Sillam-Dussés, V. Roy, C. Poteaux, R. Hanus, and Y. Roisin. 2015. Asexual queen succession in the higher termite *Embiratermes neotenicus*. Proc. R. Soc. B Biol. Sci. 282.

17. Galimov, Y., B. Walser, and C. R. Haag. 2011. Frequency and inheritance of non-male producing clones in *Daphnia magna*: Evolution towards sex specialization in a cyclical parthenogen? J. Evol. Biol. 24:1572–1583.

18. Grattapaglia, D., and R. Sederoff. 1994. Genetic linkage maps of *Eucalyptus grandis* and *Eucalyptus urophylla* using a pseudo-testcross: Mapping strategy and RAPD markers. Genetics 137:1121–1137.

19. Haenel, Q., T. G. Laurentino, M. Roesti, and D. Berner. 2018. Meta-analysis of chromosome-scale crossover rate variation in eukaryotes and its significance to evolutionary genomics. Mol. Ecol. 27:2477–2497.

20. Hebert, P. D. N. 1981. Obligate asexuality in *Daphnia*. Am. Nat. 117:784–789.

21. Hebert, P. D. N. 1978. The population biology of *Daphnia* (crustacea, Daphnidae). Biol. Rev. 53:387–426.

22. Hebert, P. D. N., M. J. Beaton, S. S. Schwartz, and D. J. Stanton. 1989. Polyphyletic origins of asexuality in *Daphnia pulex*. I. Breeding-system variation and levels of clonal diversity. Evolution 43:1004.

23. Hebert, P. D. N., and T. J. Crease. 1980. Clonal coexistence in *Daphnia pulex* (Leydig): another planktonic paradox. Science 207:1363–1365.

24. Hebert, P. D. N., and T. J. Crease. 1983. Clonal diversity in populations of *Daphnia pulex* reproducing by obligate parthenogenesis. Heredity 51:353–369.

25. Hebert, P. D. N., S. S. Schwartz, R. D. Ward, and T. L. Finston. 1993. Macrogeographic patterns of breeding system diversity in the *Daphnia pulex* group. I. Breeding systems of Canadian populations. Heredity 70:148–161.

26. Hebert, P. D. N., R. D. Ward, and L. J. Weider. 1988. Clonal-diversity patterns and breeding-system variation in *Daphnia pulex*, an asexual-sexual complex. Evolution 42:147–159.

27. Hiruta, C., C. Nishida, and S. Tochinai. 2010. Abortive meiosis in the oogenesis of parthenogenetic *Daphnia pulex*. Chromosom. Res. 18:833–840.

28. Hunter, N. 2007. Meiotic recombination. Pp. 381–442 *in* Molecular Genetics of Recombination. Springer Berlin Heidelberg, Berlin, Heidelberg.

29. Innes, D. J., and R. L. Dunbrack. 1993. Sex allocation variation in *Daphnia pulex*. J. Evol. Biol. 6:559–575.

30. Innes, D. J., C. J. Fox, and G. L. Winsor. 2000. Avoiding the cost of males in obligately asexual *Daphnia pulex* (Leydig). Proc. R. Soc. B Biol. Sci. 267:991–997.

31. Innes, D. J., and P. D. N. Hebert. 1988. The origin and genetic basis of obligate parthenogenesis in *Daphnia pulex*. Evolution 42:1024.

32. Jaquiéry, J., S. Stoeckel, C. Larose, P. Nouhaud, C. Rispe, L. Mieuzet, J. Bonhomme, F. Mahéo, F. Legeai, J. P. Gauthier, N. Prunier-Leterme, D. Tagu, and J. C. Simon. 2014. Genetic control of contagious asexuality in the pea aphid. PLoS Genet. 10:e1004838.

33. Keith, N., A. E. Tucker, C. E. Jackson, W. Sung, J. I. L. Lledó, D. R. Schrider, S. Schaack, J. L. Dudycha, M. Ackerman, A. J. Younge, J. R. Shaw, and M. Lynch. 2016. High mutational rates of large-scale duplication and deletion in *Daphnia pulex*. Genome Res. 26:60–69.

34. Kramer, M. G., and A. R. Templeton. 2001. Life-history changes that accompany the transition from sexual to parthenogenetic reproduction in *Drosophila mercatorum*. Evolution 55:748–761.

35. Lenormand, T., and J. Dutheil. 2005. Recombination difference between sexes: A role for haploid selection. Pp. 0396–0403 *in* PLoS Biology.

36. Li, H., B. Handsaker, A. Wysoker, T. Fennell, J. Ruan, N. Homer, G. Marth, G. Abecasis, and R. Durbin. 2009. The Sequence Alignment/Map format and SAMtools. Bioinformatics 25:2078–2079.

37. Li, W. H., T. Gojobori, and M. Nei. 1981. Pseudogenes as a paradigm of neutral evolution. Nature 292:237–239.

38. Lukacsovich, T., and A. S. Waldman. 1999. Suppression of intrachromosomal gene conversion in mammalian cells by small degrees of sequence divergence. Genetics 151:1559–1568.

39. Lynch, M. 1984. The limits to life history evolution in *Daphnia*. Evolution 38:465–482.

40. Lynch, M., and J. S. Conery. 2000. The evolutionary fate and consequences of duplicate genes. Science 290:1151–1155.

41. Lynch, M., A. Seyfert, B. Eads, and E. Williams. 2008. Localization of the genetic determinants of meiosis suppression in *Daphnia pulex*. Genetics 180:317–327.

42. Mather, K. 1938. Crossing-over. Biol. Rev. 13:252–292.

43. Miles, L. G., S. R. Isberg, T. C. Glenn, S. L. Lance, P. Dalzell, P. C. Thomson, and C. Moran. 2009. A genetic linkage map for the saltwater crocodile (*Crocodylus porosus*). BMC Genomics 10:1–11.

44. Moritz, R. F. A., and M. Haberl. 1994. Lack of meiotic recombination in thelytokous parthenogenesis of laying workers of *Apis mellifera capensis* (The cape honeybee). Heredity 73:98–102.

45. Normark, B. B., O. P. Judson, and N. A. Moran. 2003. Genomic signatures of ancient asexual lineages. Pp. 69–84 *in* Biological Journal of the Linnean Society.

46. Oldroyd, B. P., M. H. Allsopp, R. S. Gloag, J. Lim, L. A. Jordan, and M. Beekman. 2008. Thelytokous parthenogenesis in unmated queen honeybees (*Apis mellifera capensis*): Central fusion and high recombination rates. Genetics 180:359–366.

47. Omilian, A. R., M. E. A. Cristescu, J. L. Dudycha, and M. Lynch. 2006. Ameiotic recombination in asexual lineages of *Daphnia*. Proc. Natl. Acad. Sci. 103:18638–18643. National Academy of Sciences.

48. Paes, T. A. S. V., A. C. Rietzler, and P. M. Maia-Barbosa. 2016. Methods for selection of *Daphnia* resting eggs: the influence of manual decapsulation and sodium hypoclorite solution on hatching rates. Brazilian J. Biol. 76:1058–1063.

49. Paland, S., J. K. Colbourne, and M. Lynch. 2005. Evolutionary history of contagious asexuality in *Daphnia pulex*. Evolution 59:800–813.

50. Rastas, P. 2020. Lep-Anchor: Automated construction of linkage map anchored haploid genomes. Bioinformatics 36:2359–2364.

51. Rastas, P. 2017. Lep-MAP3: Robust linkage mapping even for low-coverage whole genome sequencing data. Bioinformatics 33:3726–3732.

52. Rebecchi, L., V. Rossi, T. Altiero, R. Bertolani, and P. Menozzi. 2003. Reproductive modes and genetic polymorphism in the tardigrade *Richtersius coronifer* (Eutardigrada, Macrobiotidae). Invertebr. Biol. 122:19–27.

53. Retnaningdyah, C., and D. Ebert. 2012. Bleach solution requirement for hatching of *Daphnia magna* resting eggs. J. Trop. Life Sci. 6:136–141.

54. Rey, O., A. Loiseau, B. Facon, J. Foucaud, J. Orivel, J. M. Cornuet, S. Robert, G. Dobigny, J. H. C. Delabie, C. D. S. F. Mariano, and A. Estoup. 2011. Meiotic recombination dramatically decreased in thelytokous queens of the little fire ant and their sexually produced workers. Mol. Biol. Evol. 28:2591–2601.

55. Schurko, A. M., and J. M. Logsdon. 2008. Using a meiosis detection toolkit to investigate ancient asexual “scandals” and the evolution of sex. BioEssays 30:579–589.

56. Schurko, A. M., J. M. Logsdon, and B. D. Eads. 2009. Meiosis genes in *Daphnia pulex* and the role of parthenogenesis in genome evolution. BMC Evol. Biol. 9:78.

57. Simon, J.-C. C., C. Rispe, and P. Sunnucks. 2002. Ecology and evolution of sex in aphids. Trends Ecol. Evol. 17:34–39.

58. Simon, J.-C., S. Stoeckel, and D. Tagu. 2010. Evolutionary and functional insights into reproductive strategies of aphids. C. R. Biol. 333:488–496.

59. Simon, J. C., F. Delmotte, C. Rispe, and T. J. Crease. 2003. Phylogenetic relationships between parthenogens and their sexual relatives: The possible routes to parthenogenesis in animals. Biol. J. Linn. Soc. 79:151–163.

60. Stapley, J., P. G. D. Feulner, S. E. Johnston, A. W. Santure, and C. M. Smadja. 2017. Variation in recombination frequency and distribution across eukaryotes: Patterns and processes. Philos. Trans. R. Soc. B Biol. Sci. 372.

61. Stelzer, C. P. 2008. Obligate asex in a rotifer and the role of sexual signals. J. Evol. Biol. 21:287–293.

62. Stelzer, C. P., J. Schmidt, A. Wiedlroither, and S. Riss. 2010. Loss of sexual reproduction and dwarfing in a small metazoan. PLoS One 5:1–6.

63. Svendsen, N., C. M. O. Reisser, M. Dukić, V. Thuillier, A. Ségard, C. Liautard-Haag, D. Fasel, E. Hürlimann, T. Lenormand, Y. Galimov, and C. R. Haag. 2015. Uncovering cryptic asexuality in *Daphnia magna* by RAD sequencing. Genetics 201:1143–1155.

64. Taylor, D. J., and P. D. N. Hebert. 1993. Cryptic intercontinental hybridization in *Daphnia* (Crustacea): The ghost of introductions past. Proc. R. Soc. B Biol. Sci. 254:163–168.

65. Tessier, A. J., and C. E. Cáceres. 2004. Differentiation in sex investment by clones and populations of *Daphnia*. Ecol. Lett. 7:695–703.

66. Tucker, A. E., M. S. Ackerman, B. D. Eads, S. Xu, and M. Lynch. 2013. Population-genomic insights into the evolutionary origin and fate of obligately asexual *Daphnia pulex*. Proc. Natl. Acad. Sci. USA 110:15740–15745.

67. van der Kooi, C. J., and T. Schwander. 2014. On the fate of sexual traits under asexuality. Biol. Rev. 89:805–819.

68. Vanin, E. F. 1985. Processed pseudogenes: characteristics and evolution. Annu. Rev. Genet. 19:253–272.

69. Veller, C., N. Kleckner, and M. A. Nowak. 2019. A rigorous measure of genome-wide genetic shuffling that takes into account crossover positions and Mendel’s second law. Proc. Natl. Acad. Sci. USA 116:1659–1668.

70. Wang, W., Z. E. Chen, and H. Li. 2021. Truncated Rank-Based Tests for Two-Part Models with Excessive Zeros and Applications to Microbiome Data. https://arxiv.org/abs/2110.05368.

71. Wersebe Matthew. 2021. The chromosome-level genome assembly and high-density genetic map of *Daphnia pulicaria* reveals the sex-specific recombination landscape of a cyclical parthenogen. Evolution 2021 online.

72. Wolinska, J., and C. M. Lively. 2008. The cost of males in *Daphnia pulex*. Oikos 117:1637–1646.

73. Xu, S., M. S. Ackerman, H. Long, L. Bright, K. Spitze, J. S. Ramsdell, W. K. Thomas, and M. Lynch. 2015a. A male-specific genetic map of the microcrustacean *Daphnia pulex* based on single-sperm whole-genome sequencing. Genetics 201:31–38.

74. Xu, S., D. J. Innes, M. Lynch, and M. E. Cristescu. 2013. The role of hybridization in the origin and spread of asexuality in *Daphnia*. Mol. Ecol. 22:4549–4561.

75. Xu, S., A. R. Omilian, and M. E. Cristescu. 2011. High rate of large-scale hemizygous deletions in asexually propagating *Daphnia*: Implications for the evolution of sex. Mol. Biol. Evol. 28:335–342.

76. Xu, S., K. Spitze, M. S. Ackerman, Z. Ye, L. Bright, N. Keith, C. E. Jackson, J. R. Shaw, and M. Lynch. 2015b. Hybridization and the origin of contagious asexuality in *Daphnia pulex*. Mol. Biol. Evol. 32:msv190.

77. Ye, Z., X. Jiang, M. E. Pfrender, and M. Lynch. 2021. Genome-wide allele-specific expression in obligately asexual *Daphnia pulex* and the implications for the genetic basis of asexuality. Genome Biol. Evol.

78. Ye, Z., C. Molinier, C. Zhao, C. R. Haag, and M. Lynch. 2019. Genetic control of male production in *Daphnia pulex*. Proc. Natl. Acad. Sci. USA 116:15602–15609.

79. Ye, Z., S. Xu, K. Spitze, J. Asselman, X. Jiang, M. S. Ackerman, J. Lopez, B. Harker, R. T. Raborn, W. K. Thomas, J. Ramsdell, M. E. Pfrender, and M. Lynch. 2017. A new reference genome assembly for the microcrustacean *Daphnia pulex*. G3 Genes, Genomes, Genet. 7:1405–1416.

