## Supplementary figures and images for "No support for a meiosis suppressor in *Daphnia pulex*: Comparison of linkage maps reveals normal recombination in males of obligate parthenogenetic lineages"

### FigureS1

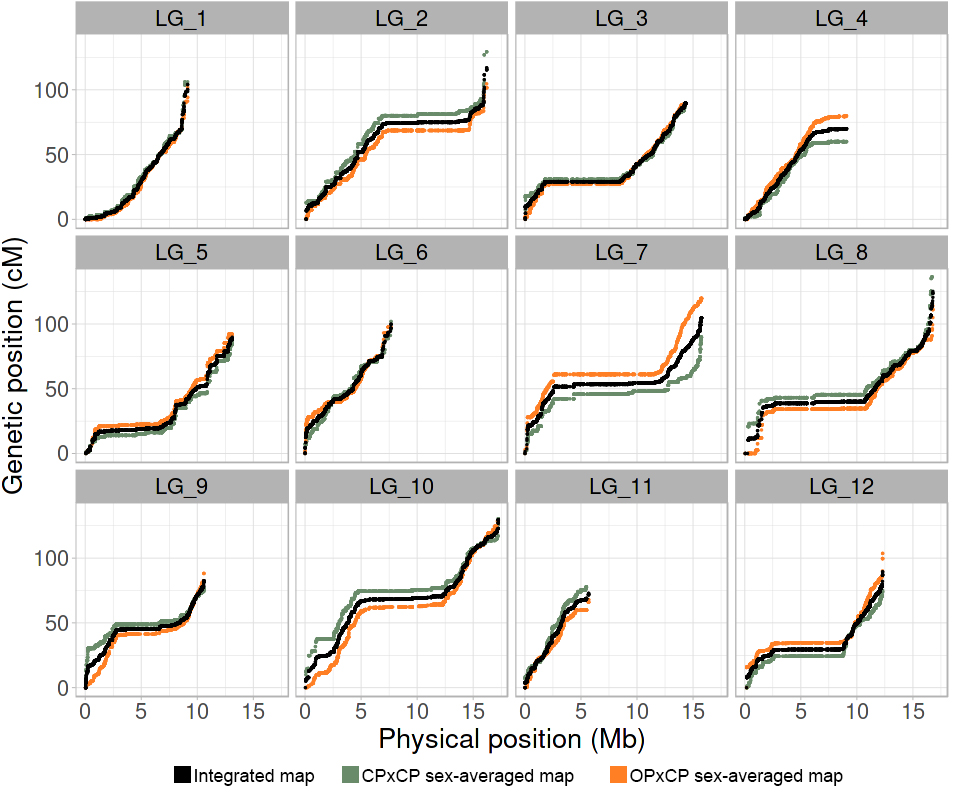
